# Cognitive modeling of the Mnemonic Similarity Task as a digital biomarker for Alzheimer’s Disease

**DOI:** 10.1101/2024.03.07.584012

**Authors:** Casey Vanderlip, Michael D. Lee, Craig E.L. Stark

**Affiliations:** Department of Neurobiology and Behavior, University of California Irvine; Department of Cognitive Science, University of California, Irvine

## Abstract

AD related pathologies, such as beta-amyloid (Aβ) and phosphorylated tau (pTau), are evident decades before any noticeable decline in memory occurs. Identifying individuals during this asymptomatic phase is crucial for timely intervention. The Mnemonic Similarity Task (MST), a modified recognition memory task, is especially relevant for early AD screening, as it assesses hippocampal integrity, a region affected (both directly and indirectly) early in the progression of the disease. Further, strong inferences on the underlying cognitive mechanisms that support performance on this task can be made using Bayesian cognitive modeling. We assessed whether analyzing MST performance using a cognitive model could detect subtle changes in cognitive function and AD biomarker status prior to overt cognitive decline. We analyzed MST data from >200 individuals (young, cognitively healthy older adults, and individuals with MCI), a subset of which also had existing CSF Aβ and pTau data. Traditional performance scores and cognitive modeling using multinomial processing trees was applied to each participants MST data using Bayesian approaches. We assessed how well each could predict age group, memory ability, MCI status, Aβ/pTau status using ROC analyses. Both approaches predicted age group membership equally, but cognitive modeling approaches exceeded traditional metrics in all other comparisons. This work establishes that cognitive modeling of the MST can detect individuals with AD prior to cognitive decline, making it a potentially useful tool for both screening and monitoring older adults during the asymptomatic phase of AD.

## 1. Introduction

Alzheimer’s disease (AD) is marked by a gradual decline in memory and cognitive abilities that are often observed only after beta-amyloid (Aβ) and phosphorylated tau (pTau) are already present (Sperling et al., 2011; Jack et al., 2018; Jia et al., 2024; Li et al., 2024). Elevated levels of Aβ and pTau increase the risk of cognitive decline (Donohue et al., 2017; Ossenkoppele et al., 2022), making this preclinical stage of AD a critical window for early detection and intervention (Sperling et al., 2013). During this phase, therapies targeting Aβ and pTau could be most effective, prior to irreversible neuronal loss (Boxer and Sperling, 2023).

Measuring Aβ and pTau is possible using PET and CSF, but both invasive and costly, limiting their general application in clinical settings (McMahon et al., 2003; Wittenberg et al., 2019). Recent developments in blood testing for Aβ and pTau levels show promise in overcoming these barriers (Hansson et al., 2023; Barthélemy et al., 2024) enabling them to become useful clinical tools. The early detection of subtle cognitive decline via digital biomarkers is also showing promise (Dagum, 2018; Ding et al., 2022; Macdougall et al., 2024). These non-invasive assessments, which can often be remotely self-administered, could complement blood tests in identifying individuals at future risk of decline, as they may detect different aspects of AD progression. Supporting this, work has found that combining blood biomarkers with cognitive tests offers a more accurate prediction of AD than using either method alone (Wang et al., 2023). However, traditional cognitive tests have been less effective in identifying individuals at high risk of AD before cognitive symptoms appear (Hedden et al., 2013). This underscores the need for refined cognitive tasks that can detect subtle cognitive changes linked to AD pathology and aid in early diagnosis when combined with biomarker analysis.

The Mnemonic Similarity Task (MST) is a promising tool as it is designed to tax hippocampal function through its emphasis on pattern separation, a process central to rapidly learning new, arbitrary information (Kirwan and Stark, 2007; Bakker et al., 2008; Lacy et al., 2011). Performance on the pattern separation component of the MST (the Lure Discrimination Index or LDI) has been associated with functional and structural changes within the hippocampus and related structures while the recognition memory aspect (REC) of the task has not (Kirwan et al., 2012; Stark et al., 2019). Given that the hippocampus (and entorhinal cortex which serves as a gateway to the hippocampus) is one of the first affected by aging and AD (Small et al., 1999, 2011; Morrison and Hof, 2002; Sabuncu, 2011), its unsurprising that performance declines with age and AD (Ally et al., 2013; Stark et al., 2013). Further, work has demonstrated that the MST can predict early cognitive changes in AD and this task has been used in multiple clinical trials including A4 and HOPE4MCI (Papp et al., 2020; Belliart-Guérin and Planche, 2023; Kim et al., 2023; Mohs et al., 2024)

The MST’s traditional metrics are designed to be simple and robust, but obscure potentially useful aspects of memory performance. Cognitive modeling of individual’s memory can give a richer understanding of mechanisms (Norman et al., 2001) and how these are altered by aging or cognitive impairments (Lee et al., 2020; Chwiesko et al., 2023; Mulhauser et al., 2023). Recently we developed a cognitive model to analyze performance on the MST using Bayesian methods that both fit individual participant performance and identified individual differences in memory and response strategies (Lee and Stark, 2023).

Here, we applied this approach to determine whether it aids the MST’s ability to discriminate various groups of individuals based on age, cognitive status, and Aβ/pTau status. We found that the cognitive model was clearly superior to traditional metrics, particularly in regards to Aβ/pTau status, highlighting the MST’s potential as an effective digital biomarker for early AD detection and monitoring.

## 2. Methods

Data from this study came from two previously published works. Experiments 1-3 used participants from Stark et al. (2013), while Experiment 4 used data from Trelle et al. (2021). Both used the same format of the MST, and both works attempted to identify cognitively “healthy” adults as part of their screening and assessment procedures.

### 2.1. Experiment 1: Predicting Age Group from Cognitive Modeling of the MST

For predicting age group, people who were less than 40 years old (n = 27, age=27.41±5.7, 16F) were classified as young and individuals who were over 60 (n = 46, age=71.33±6.4, 28F) were considered aged. All individuals were initially screened to be cognitively healthy without impairment using a battery of cognitive tasks. These include the Mini Mental State Exam (Crum et al., 1993), Wechsler Memory Scale Logical Memory (Wechsler, 1997c), Rey Auditory Verbal Learning Test (Rey, 1941), Verbal Fluency (Tombaugh, Kozak, & Rees, 1999), Digit Span (Wechsler, 1997a), Trails A and B (Tombaugh, 2004),and Letter Number Sequencing (Wechsler, 1997b), and the Wechsler Adult Intelligence Score III (Wechsler, 1997a). All individuals scored within 1.5 standard deviations of the mean of their age group for all neuropsychological measures.

### 2.2. Experiment 2: Predicting memory deficits older adults using Cognitive Modeling of the MST

Significant work has used the Rey Auditory Verbal Learning Test (RAVLT) to differentiate older adults into separate groups based on cognitive function. The RAVLT consists of learning a list of 15 words and recalling them after a delay of 15 minutes and the delay score ranges from 0 to 15 and reflects the number of words correctly recalled after the delay. In the original report, older adults were split into thirds based on their RAVLT performance to parallel work in the rodents that examined aged unimpaired (AU) and aged impaired (AI) groups (Stark et al., 2013). It is important to note that AI individuals (RAVLT of 5-8) are still within their age-based norms and are not clinically impaired. AU individuals (RAVLT of 12-15) have performance similar to young adults (this threshold is often used as part of the “SuperAger” criteria). However, here we used a threshold of 9 on the RAVLT to split older adults into either individuals with age-related memory deficits (AMD) or no age-related memory deficits (NMD). Similar to prior work, individuals who scored higher than 9 were considered NMD (n = 31, age=71.29±6.79, 18F), and those who scored 9 or below, but within normal limits of their age group, were considered AMD (n = 15, age=71.40±5.8, 10F) (Harrison et al., 2012; Gefen et al., 2014, 2015; Radhakrishnan et al., 2022).

### 2.3. Experiment 3: Predicting cognitive status in cognitively older adults

To predict whether older adults were cognitively normal (CN) or had mild cognitive impairment (MCI) using the MST, the same 46 adults over the age of 60 from experiments 1 and 2 were used for older adults who are cognitively intact (n = 46, age=71.33±6.4, 28F). A further 10 individuals (age=76.30±6.78, 5F) who were diagnosed with amnestic MCI were also included. Individuals with MCI were diagnosed by the UCI Alzheimer’s Disease Research Center (ADRC). All individuals with MCI had a CDR global rating of 0.5, a memory complaint and impaired memory function on neuropsychological testing. Final diagnosis of amnestic MCI was reached by neurologists and neuropsychologists at clinical consensus conferences within the UCI ADRC. All participants had no history of neurological or psychiatric disorders, head trauma with loss of consciousness, drug abuse or dependency.

### 2.4. Experiment 4: Predicting biomarkers of Alzheimer’s disease in cognitively normal older adults

Experiment 4 used previously published data (Trelle et al., 2021), collected as part of the Stanford Aging and Memory Study (SAMS). 133 older adults (age = 68.8±5.8, 83F) were administered the MST and underwent a lumbar puncture to quantify AD biomarkers. All individuals had normal or corrected-to-normal vision/hearing, were right-handedness, were native English speakers, and no history of neurologic or psychiatric disease. Further, each participant had a Clinical Dementia Rating (CDR) global score of zero and performance within the normal range on a standardized neuropsychological test battery. Lastly, all participants were deemed cognitively normal during a clinical consensus meeting consisting of neurologists and neuropsychologists. The previously derived Aβ42, Aβ40, and p-tau181 levels were used in the present analyses (see Trelle et al., 2021 for details).

### 2.5. Mnemonic Similarity task

The MST is a widely used cognitive task that is thought to critically tax hippocampal pattern separation (Fig 1A; Stark et al., 2013, 2019). Both data sources used the traditional version of the MST, which consists of an incidental encoding phase and an explicit test phase. During the encoding phase, individuals made successive indoor/outdoor judgments for 128 images (2s each, 0.5 ISI, color objects on a white background) via a button press. Immediately following the encoding phase, participants were given instructions for a recognition memory test where they were told to identify objects as either “Old” (the exact same picture as before), “Similar” (indicating this is similar to, but not identical to the studied item – e.g., a different exemplar, a rotation, etc.), or “New” via a button press. Here, participants saw 192 images (2s each, 0.5 ISI) and responded to each of these images. Images consisted of 64 exact repeats from the encoding phase (targets), 64 completely novel images (foils) and 64 images that were similar, but not identical to images seen during encoding (lures).

**Figure 1.**
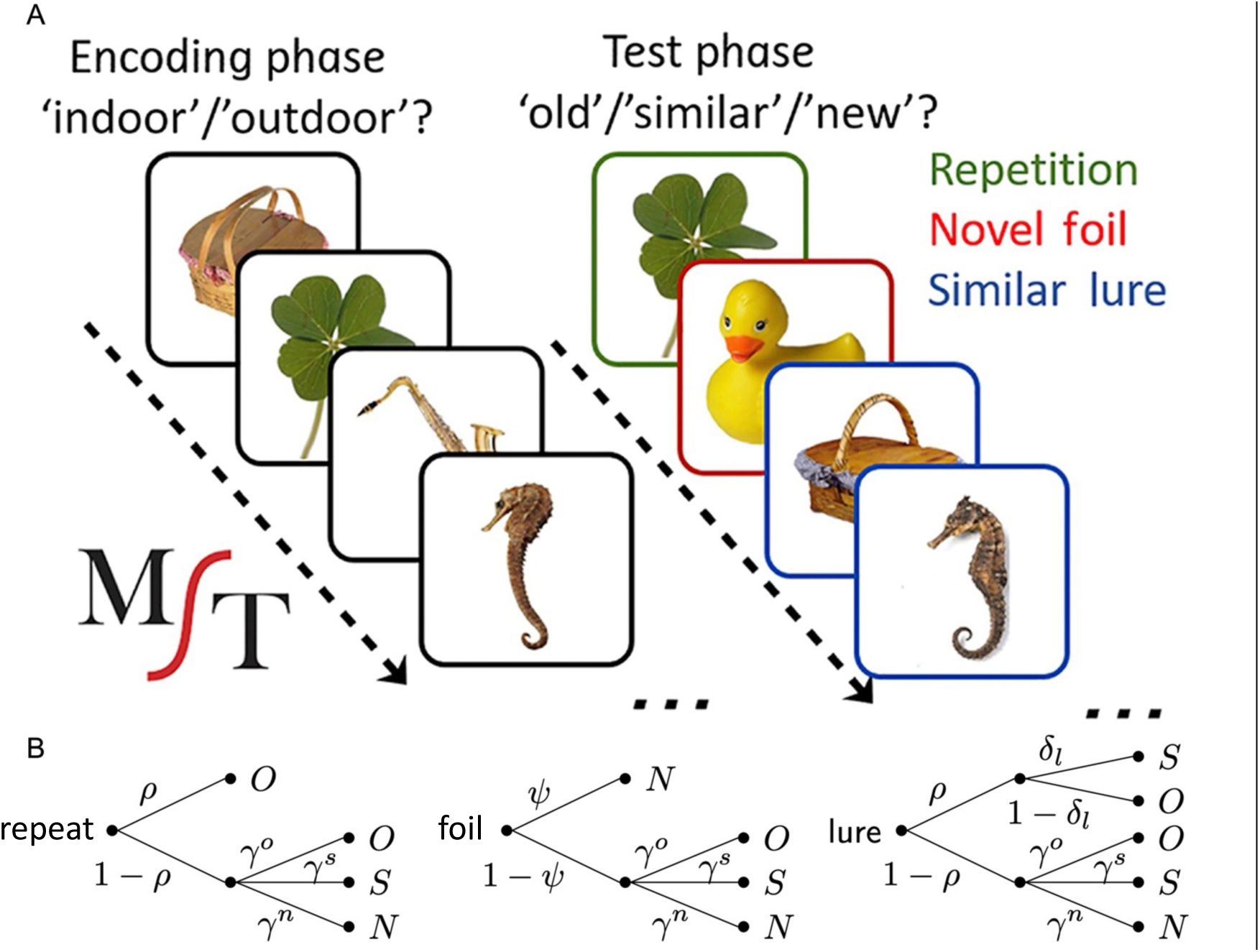
Cognitive modeling of the MST. A) Sample stimuli used during the incidental encoding phase and the subsequent Old/Similar/New recognition task B) Probability tree diagrams of the MPT model, demonstrating the decision-making process utilized within the Old/Similar/New version of the MST.

Multiple behavioral metrics were extracted from the MST (Table 1), including the traditional recognition memory (REC) and Lure Discrimination Index (LDI) scores. REC is a commonly used measure of recognition memory and is the probability of “Old” responses given to the target items minus the corresponding probability of “Old” responses given to the foils (to correct for response bias). To quantify ability to discriminate between similar lures, the LDI is the difference between the probability of giving a “Similar” response to lure items and the probability of giving a “Similar” response to the foils to account for any bias individuals may have in using the “Similar” response overall. For a follow-up analysis, we also quantified the rate of “Old” responses for target trials (hits), rate of “Similar” responses for lure trials (correct rejections of lures) and rate of “New” responses for foil trials (correct rejections of foils). Further, we attempted to get a readout of guessing by calculating the rate of “Old” responses on foil trials, the rate of “Similar” responses on foil trials, and the rate of “New” responses on target trials.

**Table 1.**

| Metrics | Type | Definition |
| --- | --- | --- |
| REC | Traditional | Recognition memory score |
| LDI | Traditional | Reflects ability to discriminate between similar lures |
| p(Old\|Repeat) | Traditional | Probability of responding old for repeats |
| p(Sim\|Lure) | Traditional | Probability of responding similar for lures |
| p(New\|Foil) | Traditional | Probability of responding new for foils |
| p(Old\|Foil) | Traditional | Probability of responding old for foils |
| p(Sim\|Foil) | Traditional | Probability of responding similar for foils |
| p(New\|Repeat) | Traditional | Probability of responding new for repeats |
| $\rho$ | Modeled | probability of remembering items at a gist level |
| $\lambda$ | Modeled | ability to discriminate remembered items from lures |
| $\psi$ | Modeled | probability of remembering that an item was not studied |
| $\gamma^o$ | Modeled | probability of guessing old |
| $\gamma^s$ | Modeled | probability of guessing similar |
| $\gamma^N$ | Modeled | probability of guessing new |

### 2.6. Cognitive modeling

Cognitive modeling provides a useful tool for inferring latent psychological variables beyond traditional measurements. Previously, we used cognitive modeling to model subject-level performance on the MST in young adults (Lee and Stark, 2023) using the multinomial processing tree (MPT) framework, a common approach for cognitive modeling of recognition memory tasks. The MPT framework assumes that cognitive processes can be divided into discrete categories or decision points (Fig 1B). Briefly, when a repeated item appears, we assume there is a probability (π) that the item is successfully matched with memory in at least a basic gist or “familiarity” form, leading to an “old” response. Failing that, we assume that guess is made with unique probabilities (response biases) for each of the three responses. Similarly, when an unrelated foil is present, there is a probability (ψ) that that the lack of a match to memory is sufficiently clear that a “no” response is made and, failing that, a three-choice guess is made.

When a similar lure is presented, there is an initial decision point involving recognizing some degree of match between the object presented and the memory of one previously encountered, based on the same ρ as above. This level of match is modeled to reflect a simpler, item-, gist-, or familiarity-based match (for both lures and repeated items). If this is unsuccessful a 3-choice guess happens as before. If successful, there is a second decision point based on a set of similarity-based probabilities (δ) capturing whether the memory retrieval contains the richer details required to reject the item as only being similar to the studied item. If successful, a “similar” response is made and if unsuccessful, an “old” response is made.

Posterior distributions for metrics within MPT models were estimated at the subject level from trial-by-trial data experimental data using JAGS. We used posterior means as point estimates for multiple metrics of interest (Table 1). These metrics include ρ, which reflects the probability of remembering items, λ, based on δ and designed to capture the ability to discriminate remembered items from lures, ψ, the probability of remembering that an item was not studied, γ^O^ (probability of guessing old), γ^N^ (probability of guessing new) and γ^S^ (probability of guessing similar).

### 2.7. Statistical analyses

All analyses were done in Python. Logistic regressions were run using statsmodels (Seabold and Perktold, 2010) to predict age group, clinical status, biomarker status, etc. from various sets of metrics. Areas under the curve (AUC) measures were derived from ROC curves of the logistic regressions. To compare model fits, we calculated the Bayesian Information Criteria (BIC) of each model (Raftery, 1995). Absolute differences in BICs of greater than 2 were considered reliable. Importantly, the logistic regressions differed in the number of variables used as predictors and it is reasonable to assume that there will be shared variance between model based and traditional metrics. Therefore, to identify how each variable acts in conjunction with the others, we performed an 8-choose-4 combinatorial analysis and quantified the number of times each metric appeared in the top third of AUCs from 8-choose-4 analyses. Independent sample t-tests were used to examine group differences in traditional and model-based metrics (Student, 1908). To investigate group changes in guessing strategies, Kolmogorov–Smirnov tests were used because data was proportioned and therefore not normally distributed. For all analyses, p < 0.05 was considered reliable.

## 3. Results

Previously, we demonstrated that the traditional REC correlated with ρ, while LDI correlated with λ (Lee and Stark, 2023, previously denoted as τ). Our first goal was to assess the relationship among the traditional and modeled metrics in the two datasets (Stark et al., 2013; Trelle et al., 2021). Like the prior work, we found strong correlations between these variables in in both datasets (Stark et al., 2013; REC vs ρ: r = 0.73, LDI vs λ: r= − 0.90, Trelle et al., 2021; REC vs ρ: r = 0.77, LDI vs λ: r = − 0.90). These results demonstrate that the model-based metrics derived here are similar to prior findings. With this, we conducted four separate experiments to assess if traditional or model-based metrics were superior in identifying individuals at risk for AD.

### 3.1. Experiment 1: Traditional metrics and model-based metrics of the MST equally predict age group status

Given that extensive work has demonstrated that older adults are impaired on the MST, we assessed whether cognitive modeling could enhance the ability to differentiate younger and older adults (Stark et al., 2013, 2019). Considered individually, There was no reliable difference in REC between age groups, while LDI was significantly lower in older adults (REC: t(71) = 1.19, p = 0.28, LDI: t(71) = 5.71, p < 0.0001). When examining modeled metrics individually, ρ showed no reliable age differences, while γ and were lower in older compared to younger adults (ρ: t(71) = 0.22, p = 0.83 ψ: t(71) = 2.62, p < 0.05,: t(71) = −5.69, p < 0.0001). A multiple logistic regression using the traditional LDI and REC as predictors achieved an AUC of 0.86 (Fig. 2A, p < 0.0001). Model-based metrics, with ρ, ψ,τ, along with guessing strategies (γ^O^ γ^N^ and γ^S^) as predictors, yielded a similar AUC of 0.84 (Fig. 2A, top, p < 0.0001), suggesting that model-based metrics did not outperform traditional metrics in predicting age group.

**Figure 2.**
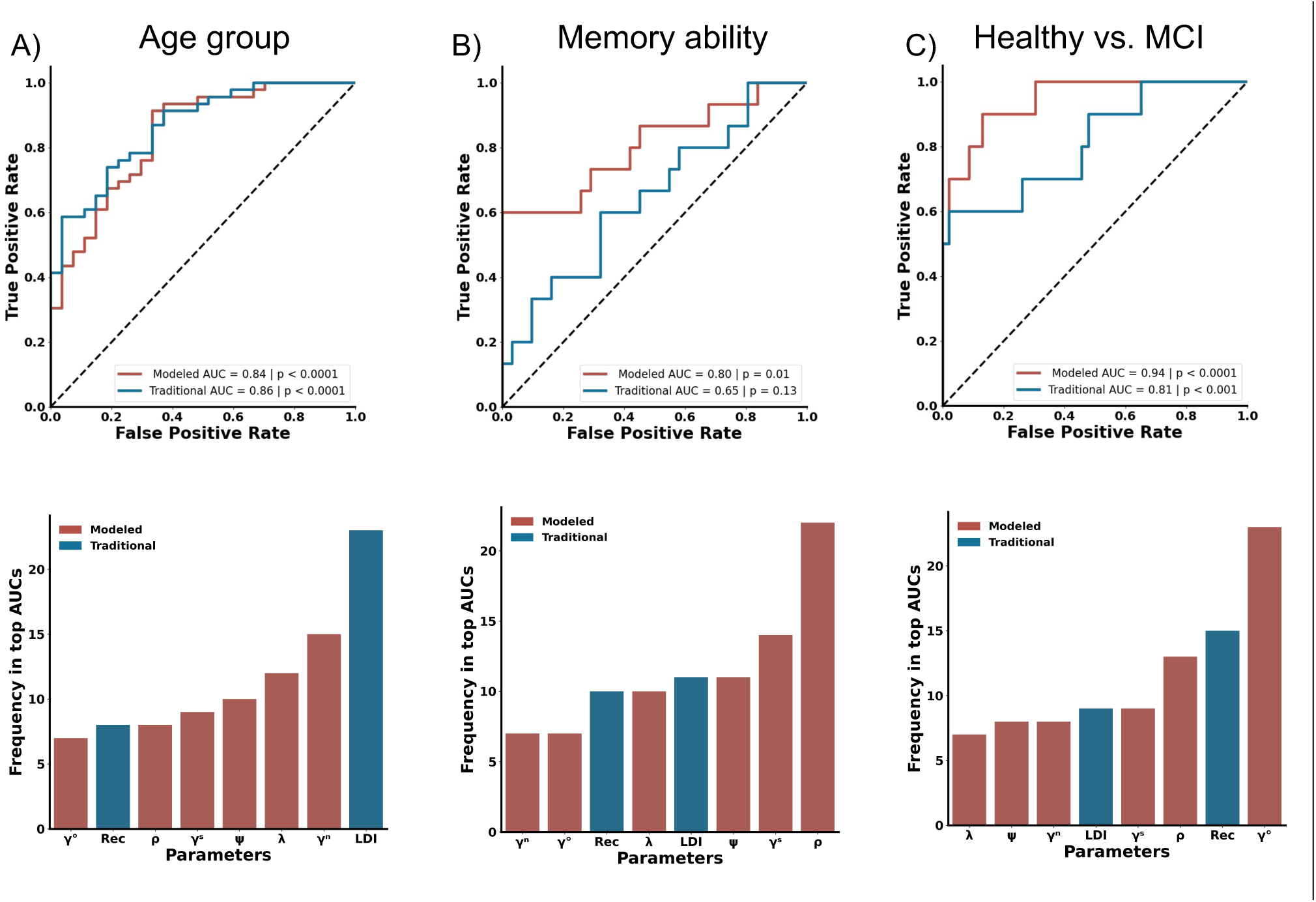
Comparison of traditional (blue) to cognitive modeling (red) performance in Experiments 1-3. ROC curves (top) frequency of presence in the top-30 AUCs in the n-choose-4 combinatorial analyses (bottom) are shown. A) Comparison of age group predictions showing no significant difference between traditional measures and cognitive model-based measures. LDI emerges as the most frequent metric in the top third of AUCs. B) Metrics derived from cognitive modeling better predict performance high-vs. low-performing older adults. The gist-based recognition memory signal metric (ρ) is the predominant metric in the n-choose-four analysis for predictive accuracy. C) Cognitive modeling metrics were superior at identifying healthy vs. MCI. Within the n-choose-four analysis, γ^O^ is the leading metrics for MCI prediction.

Considering metrics in isolation and considering them in combination with other metrics from the same approach does allow for direct comparisons across the techniques. However, as shown above, the metrics are not independent of each other, and the two approaches differ in the number of variables considered. To appreciate better the impact each variable might have in conjunction with the others, we performed an 8-choose-4 combinatorial analysis and identified how often each factor occurred in the top third of resulting AUCs. This revealed that the LDI was the most common metric in distinguishing younger and older adults, appearing in virtually all the top-performing models and almost twice as often as the most frequent cognitive model-based metric (Fig. 2A, bottom). Thus, when considering the simpler task of predicting age group membership, we found no evidence that cognitive modeling was superior to the traditional approach.

### 3.2. Experiment 2: Model-based metrics better identify memory ability older adults

Differing cognitive ability in older adults can be informative of future decline. Therefore, we next asked if performance on the MST along with cognitive modeling could aid in dissociating across levels of cognitive function in healthy adults by discriminating NMD versus AMD. Considering each variable individually, REC and LDI levels were similar in NMD and AMD (REC: t(44) = 0.22, p = 0.22, LDI: t(44) = 1.72, p = 0.09). When measuring model-based metrics, ρ was significantly higher in NMD compared to AMD individuals with no difference in ψ or λ(ρ: t(44) = 3.10, p < 0.01; ψ: t(44) = 1.248, p = 0.22;: λ t(44) = −0.31, p = 0.76). When combining LDI and REC, a multiple logistic regression did not successfully distinguish NMD versus AMD (AUC = 0.65, p = 0.13, Figure 2B). However, a multiple logistic regression with model-based metrics were able to stratify NMD from AMD with an AUC of 0.80 (p < 0.05). When assessing combinations of traditional and model-based metrics in an 8-choose-4 combinatorial analysis, ρ emerged as the most consistent metric in the top-performing models with LDI appearing as a distant 4^th^ most consistent (Fig. 2B, bottom). This suggests that cognitive modeling provides a more accurate identification of memory ability in older adults than traditional metrics, but that this is driven heavily by the model’s estimate of how well individuals remember at least the gist of an item.

### 3.3. Experiment 3: Model-based metrics better predict MCI status

We next investigated whether cognitive modeling of the MST could better identify individuals with MCI compared to traditional metrics. We found that individuals with MCI had significantly lower REC performance compared to cognitively normal older adults, but there were no differences between groups in LDI scores (REC: t(52) = 4.73, p < 0.0001; LDI: t(52) = 0.77, p = 0.44). We also found that ρ decreased in individuals with MCI, but no difference in groups for ψ or λ (ρ: t(52) = 5.51, p < 0.0001; ψ: t(52) = 0.99, p = 0.33; λ: t(52) = −0.16, p = 0.87). In the multiple logistic regression, we found that the combination of REC and LDI could classify MCI status with good accuracy (AUC = 0.81, p < 0.001, Figure 2C). However, cognitive model-based metrics offered superior predictive power, achieving an AUC of 0.94 (p < 0.0001). Permutation analysis found that γ^O^ was the most influential metric, appearing in all the top third of models (Fig. 2C, bottom). This suggests cognitive modeling is superior at detecting MCI over traditional metrics largely due to the ability to derive differences in guessing strategy on the task.

### 3.4. Experiment 4: Model-based metrics can better predict Aβ and Tau status in cognitively normal older adults

We next evaluated whether cognitive modeling of the MST could detect Aβ status in cognitively healthy older adults, classified as Aβ+ or Aβvia CSF Aβ42/Aβ40 ratios. Aβ+ individuals had decreased REC scores but equivalent LDI performance compared to Aβcounterparts (REC: t(131) = 2.68, p < 0.01; LDI: t(131) = 0.33, p = 0.74). Further, ρ was lower in Aβ+ compared to Aβ-older adults with no group differences in χ and λ (ρ: t(131) = 2.54, p < 0.05; λφ./: t(131) = 1.11, p = 0.27;: t(131) = −0.53, p = 0.60). A multiple logistic regression with traditional metrics could modestly predict amyloid status (AUC = 0.64, p < 0.05, Fig. 3A). On the other hand, a multiple logistic regression with model-based metrics better predicted amyloid status (AUC = 0.73, p < 0.05). When conducting an 8-choose-4 combinatorial analysis to investigate the impact each variable might have in relation with the others, γ^O^ was the most predictive metric among the top third of AUCs. Interestingly, γ^O^ was represented in nearly all the top models and twice as often as both traditional metrics (Fig. 3A, bottom). Cognitive modeling thus better identifies asymptomatic individuals with elevated amyloid burden due to its ability to derive differences in guessing old.

**Figure 3.**
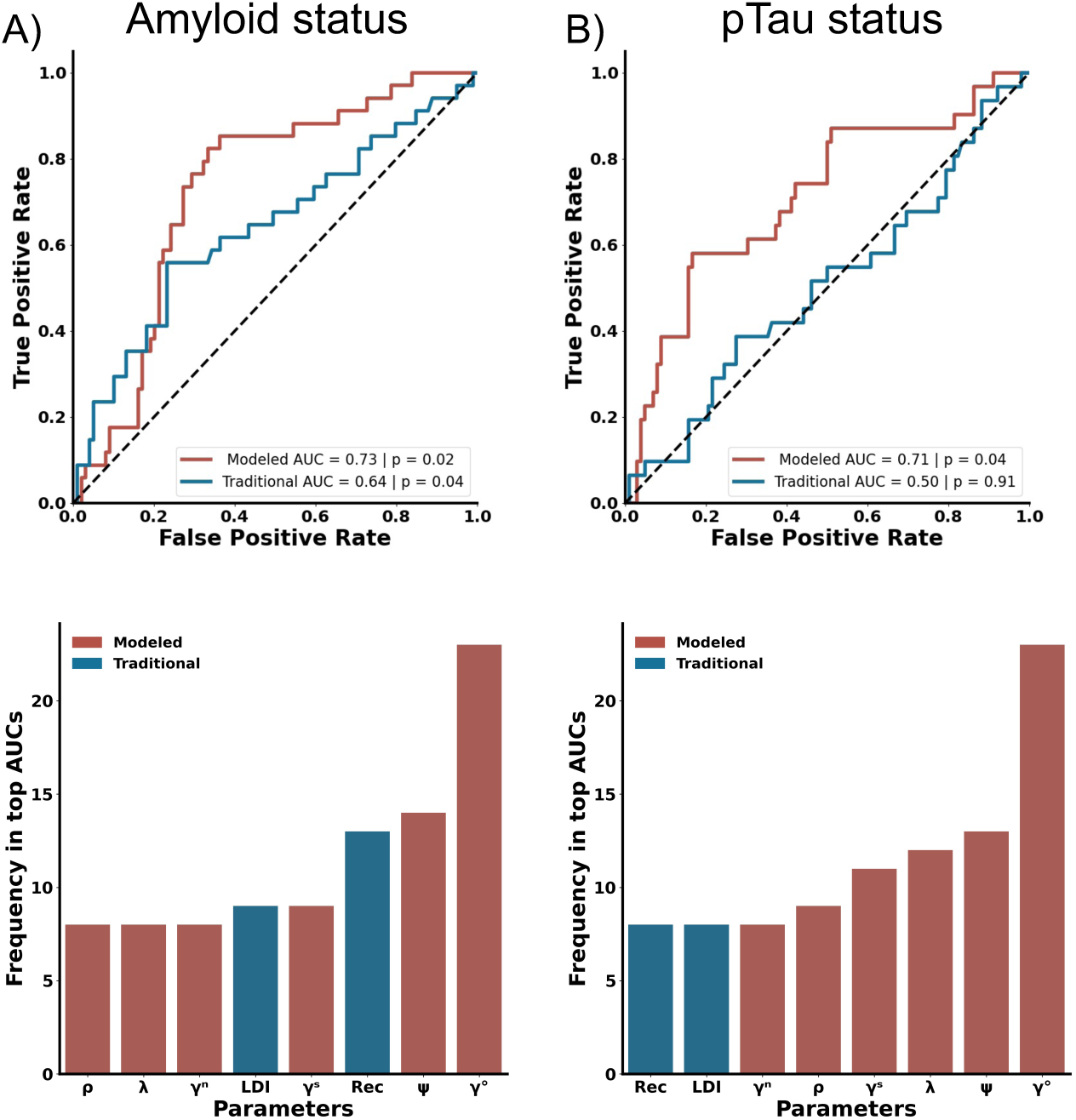
Utilization of Cognitive Modeling in the MST for Predicting AD Biomarker Status. A) Both traditional and cognitive modeling-derived metrics can predict Aβ status with cognitive model-based metrics showing superior predictive accuracy. The metric γ^O^ is identified as the most frequently occurring metric in the top results from n-choose-four analyses for predicting Aβ status. C) Cognitive modeling, but not traditional measures can successfully predict pTau status. D) The metric γ^O^ is again highlighted as the most common metric in the top 30 AUCs from a n-choose-four analysis for predicting pTau status. For panels A-B, Traditional measures are in blue, while cognitive model-based metrics are in red.

While both Aβ and pTau are biomarkers for AD, pTau has a stronger link to cognitive decline and may better predict disease progression. Somewhat surprisingly, cognitively normal older adults with elevated pTau levels did not differ on either traditional or model-based metrics (REC, LDI, ρ, γ and) compared to those with normal pTau levels (all ps > 0.10). Likewise, a multiple logistic regression with REC and LDI failed to predict pTau status (AUC = 0.50, p = 0.91, Fig. 3B). Importantly, the logistic regression with the model-based metrics did predict pTau status (AUC = 0.71, p < 0.05). When conducting an 8-choose-4 combinatorial analysis, the single clearly most reliable metric was γ^O^, appearing more than twice as much as the next most important metric (γ) (Fig. 3B, bottom). Further, every metric from cognitive modeling were more represented than REC and LDI in the top third of models. Overall, cognitive modeling outperformed traditional metrics in predictive accuracy, suggesting its effectiveness in early AD screening.

### 3.5. Changes in model-derived guessing strategies with age, cognitive impairment, and biomarker status

Given that modeling guessing probabilities were informative for predicting AD biomarker status and cognitive status, we took a deeper dive into guessing strategies. We first explored how model-based guessing strategies change across age groups. Cognitive modeling suggested that younger adults tended to guess “similar” more frequently and “new” less frequently than older adults (Kolmogorov–Smirnov test; γ^S^: D = 0.37, p < 0.05; γ^N^: D = 0.38, p < 0.05), without any significant age-related differences for guessing “old” (γ^O^: D = 0.25, p = 0.20). This pattern suggests an age-related shift from guessing “similar” to “new.” Interestingly, however, no significant differences in guessing strategies were found between healthy older adults with and without memory deficits (Kolmogorov–Smirnov test: all p > 0.10). In contrast, older adults with MCI were more inclined to guess “old” and less likely to guess “similar” or “new” compared to cognitively healthy older adults (Kolmogorov–Smirnov test; γ^O^: D = 0.6318, p < 0.01; γ^N^: D = 0.4910, p < 0.05; γ^S^: D = 0.5318, p < 0.05). These results underscore that aging and MCI distinctly affect guessing strategies.

We next investigated how guessing strategies on the MST varied with AD biomarker status in older adults. Cognitive modeling suggested that those with elevated amyloid displayed a tendency to guess “old” more frequently than their counterparts without elevated amyloid, but this failed to reach significance, Further, there were no significant differences in biases towards guessing “similar” or “new” (Kolmogorov–Smirnov test; γ^O^: D = 0.24, p = 0.08, γ^S^: D = 0.21, p = 0.18, γ^N^: D = 0.13, p = 0.75). We next investigated whether guessing strategies changed as a function of pTau status. We observed that individuals with elevated pTau levels were more likely to guess “old” and less likely to guess “similar,” with no change in the likelihood of guessing “new” (Kolmogorov–Smirnov test; γ^O^: D = 0.32, p < 0.05; γ^S^: D = 0.34, p < 0.05, γ^N^: D = 0.19, p = 0.34). These differences in predicted guessing strategies further highlight the benefits of cognitive modeling of the MST.

### 3.6. Addition of raw metrics of guessing do not match model-based metrics

Given the differences in guessing across performance ability, impairment level, amyloid and pTau status in older adults, we next asked whether we could derive guessing strategies based on response patterns. Specifically, we used the proportion of trials an individual responded “old” on foils as a measure of guessing old, the proportion of trials an individual responded ‘similar’ on foils as a measure of guessing similar, and the proportion of trials an individual responded ‘new on repeats as a measure of guessing new. REC and LDI incorporate these metrics in their calculations as part of their difference scores, intended to factor out differences in guessing rates. Any baseline shift in the probability of guessing “old” or “similar” would presumably affect both components of the difference metrics, removing what the model-based analyses suggest could be highly informative. Therefore, we also added the raw proportion of trials people responded New for Foil trials, the proportion of trials an individual responded ‘old’ on repeats and the proportion of trials an individual responded ‘similar’ on lure trials. Using these six new metrics, we asked whether these metrics could increase the predictive value of the MST to the same level as that provided by cognitive modeling.

We first asked whether these new behavioral metrics could predict whether cognitively normal older adults exhibited memory deficits. In a multiple logistic regression, we found that while the raw AUC appeared elevated, this model could still not reliably predict performance level (AUC = 0.78, P = 0.21, Fig. 5A). We also investigated whether these differing models better fit the data by examining their respective BIC which is a metric that reflects goodness-of-fit of a model (Wagenmakers and Farrell, 2004). Using this measure, we found that the cognitive model-based metrics better fit the data compared to the probability based traditional metrics (Cognitive modeling BIC: 76.49, Raw traditional BIC: 81.86) suggesting that, despite the addition of the raw traditional metrics, cognitive modeling better predicts memory ability. Next, when assessing whether these new probability-based metrics helped with prediction of MCI status, we found that a multiple logistic regression with these raw traditional metrics significantly bolstered the predictive accuracy of the MST yielding an AUC of 0.95 (p < 0.0001, Fig. 5B), which aligns with the predictive strength of cognitive modeling and actually provided a better fit of the data than model-based metrics (Cognitive model-based BIC: 49.77, Raw traditional BIC: 45.12).

**Figure 4.**
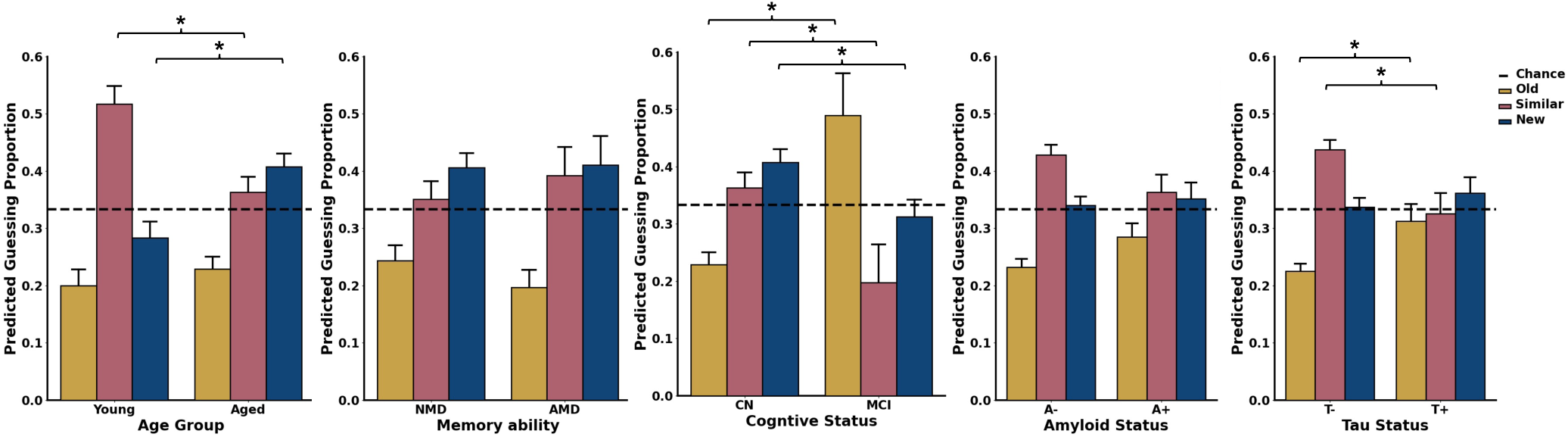
Influence of Age, memory ability, cognitive status, and AD Biomarker Status on modeled guessing strategies in the MST. A) Compared to younger adults, older adults exhibit a higher likelihood of guessing new and a lower tendency to guess similar. B) Among older adults, those with no agerelated memory deficits show no significant differences in guessing strategies when compared to individuals with age-related memory deficits. C) Individuals with MCI demonstrate a greater bias towards guessing old and are less inclined to guess similar or new relative to cognitively healthy older adults. D) There are no differences in guessing strategies between Aβ- and Aβ+ older adults. E) Older adults with elevated pTau levels display a marked shift from guessing similar towards guessing old compared to pTau negative older adults. For panels A-E, the metric for guessing old (γ^O^) is represented in yellow, for guessing similar (γ^S^) in pink, and for guessing new (γ^N^) in blue.

**Figure 5.**
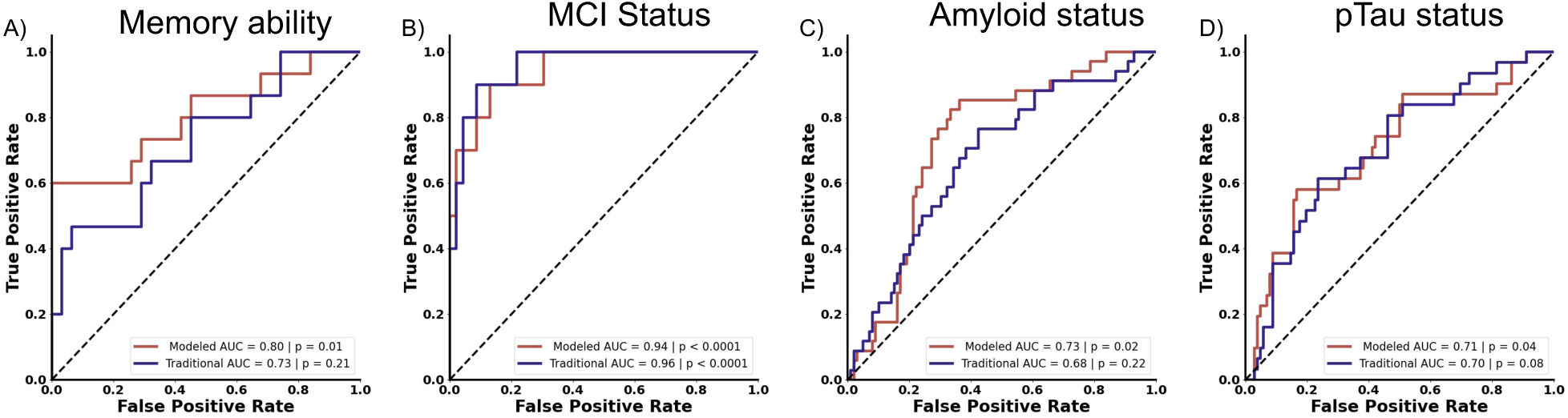
Additional raw-derived measures may improve upon traditional metrics, but do not match the cognitive model-based metrics. A) Adding raw metrics of guessing fail to predict memory ability, in contrast to cognitive modeling measures. B) In cases of overt cognitive impairment, such as MCI, raw-derived guessing measures show comparable predictive value to that of cognitive modeling. C) Additional raw metrics fail to match cognitive modeling in predicting Aβ status. D) While the additional raw measures improve the predictive ability of traditional approaches for pTau status, their performance does not exceed chance levels. For panels A-D, traditional measures are depicted in purple, and cognitive model-based metrics are illustrated in red.

We next evaluated whether raw traditional metrics could match cognitive modeling in predicting amyloid status. A multiple logistic regression with these metrics did not match the predictive capacity of model-based metrics (AUC = 0.68, p = 0.22, Fig. 5C). Moreover, when comparing model fits using the BICs, model-based metrics better fit the data (Cognitive modeling BIC: 167.18, Raw traditional BIC: 177.12). We next asked if the new probability-based metric could predict pTau status. These new metrics did show a qualitative improvement in predicting pTau status, but this was not statistically reliable (AUC = 0.70, p = 0.08, Fig. 5D) Further, model-based metrics better fit the data compared to probability-based metrics (Cognitive modeling BIC: 161.95, Raw traditional BIC: 167.52), reinforcing the superiority of cognitive modeling in predicting amyloid and pTau status.

## 4. Discussion

The MST is a widely-used memory test that assesses changes in hippocampal integrity in various conditions including age-related cognitive decline and Alzheimer’s disease (Stark et al., 2019). Given that this task is resistant to practice effects and can be easily performed remotely, it has emerged as an ideal candidate for clinical use as a digital biomarker for AD. However, work investigating whether performance on the MST exceeds traditional neuropsychological tests in stratifying individuals with and without cognitive impairment has yielded mixed results (Belliart-Guérin and Planche, 2023; Kim et al., 2023). We have previously demonstrated that cognitive modeling can be applied on the MST and shown how multiple cognitive metrics can be inferred from these models, but we did not know whether these new metrics would aid the predictive value of the MST. In this study, we used data from multiple studies of aging, MCI, and AD biomarkers, to compare the predictive value of traditional metrics versus model-based metrics. We found that cognitive modeling enhances the ability of the MST in identifying older adults at risk of developing AD prior to cognitive decline. This work demonstrates that the MST is well-suited to enhance early diagnosis in AD thereby enabling earlier intervention strategies to treat this disease.

### 4.1. Cognitive Modeling of the MST identifies differing cognitive capacity in older adults

A large body of work has demonstrated that advancing age is associated with significant impairment on the LDI metric of the MST, while REC remains stable with age (Yassa et al., 2010; Stark et al., 2013; Gellersen et al., 2021). Similarly, we found that traditional behavioral metrics of the MST predicted age group with high proficiency (AUC = 0.85) and cognitive modeling did not increase the high predictive value of the MST. Further analyses showed that the most important metric is LDI, appearing in nearly all top performing models in a permutation analysis and almost twice as much as all other measures. This further reaffirms that age-related impairments on the MST are due, in part, to deficits in pattern separation and is consistent with a hippocampal contribution to age-related impairments.

While age-related impairments are seen on many cognitive tests, there is typically significant heterogeneity within the aging population with a subset of healthy older adults exhibiting age-related memory deficits while others show young-like performance (e.g., “SuperAgers” or “Aged unimpaired). Our analyses showed that traditional measures of MST performance did not readily distinguish these two, but that the cognitive modeling approach could. Notably, the most predictive metric from our cognitive modeling was ρ, a metric indicative of memory retention that we hypothesize to be analogous to REC. Intriguingly, unlike ρ, REC did not differentiate between older adults with and without memory deficits, suggesting that ρ may be a more nuanced and sensitive measure of subtle memory differences. Critically, future work will be needed to understand the neural mechanisms that account for these changes in memory capacity.

We next explored the potential of the MST to identify individuals with MCI, often considered a precursor to or risk factor for AD and other dementias. The diagnosis of MCI is frequently missed, with perhaps only ∼8% of those affected accurately identified (Mattke et al., 2023; Liu et al., 2023). Closing this diagnostic gap is therefore crucial. We demonstrated that, like previous work, performance on the MST is a reliable predictor MCI with an AUC of 0.81 (Kim et al., 2023, Belliart-Guérin et al., 2023). However, the predictive accuracy was significantly improved to an AUC of 0.94 when cognitive modeling techniques were applied. This substantial increase in discriminative power suggests that cognitive modeling of the MST has the potential to be an effective tool in clinical practice, enhancing the identification of cognitive impairment and potentially mitigating the current underdiagnosis of MCI.

### 4.2. Cognitive Modeling of the MST predicts AD biomarker status in cognitively healthy older adults

Recent studies have utilized comprehensive cognitive batteries to identify individuals at higher risk of AD (Lim et al., 2016; Papp et al., 2020; Macdougall et al., 2024), with longitudinal cognitive testing used to identify healthy adults with elevated Aβ and pTau levels (Lim et al., 2016; Jutten et al., 2022; Papp et al., 2023). Critically, the cognitive battery used in this study included a shortened version of the MST (there, called the BPSO) and longitudinal changes on the MST could better predict memory impairment over three months compared to baseline neuropsychological scores. Importantly, none of the other tasks within the cognitive battery could exceed baseline neuropsychological scores. Despite these advancements, the ideal cognitive test for AD would be one that is quick, easily accessible, and capable of being completed in a single clinic visit or at home, all while reliably predicting AD biomarker status. The development of such a task could significantly enhance early diagnosis and intervention strategies for Alzheimer’s disease.

Cerebral Aβ deposition is present up to 20 years before clinical cognitive symptoms are detected, highlighting the need for more sensitive tasks (Sperling et al., 2011; Li et al., 2024). A recent study demonstrated that combining performance on multiple versions of the MST could modestly predict amyloid status in cognitively normal older adults (Kim et al., 2023). We found that, like this work, performance on the MST could modestly predict Aβ status (AUC=0.64).

Critically, however, cognitive modeling enhanced the ability of the MST to predict Aβ status reaching an AUC of 0.73. Together, our results reaffirm that performance on the MST is related to Aβ status and extends prior findings by demonstrating that the addition of inferred cognitive mechanisms enhances the predictive value of the MST.

Aβ buildup is known to drive pTau accumulation, yet it is pTau that exhibits a stronger connection to cognitive decline and that amyloid accumulation, in the absence of pTau, does not correlate with cognitive impairment (Desikan et al., 2012). Conversely, pTau has been implicated in hippocampal hyperactivity, widespread neurodegeneration, and the transition to dementia (Berron et al., 2019; Ossenkoppele et al., 2022). Further, a prior study demonstrated that elevated pTau within the medial temporal lobe correlates with impairments in pattern separation (Maass et al., 2019). However, it is not known if increased pTau levels can be predicted from performance on the MST. Interestingly, MST performance alone was not able to predict elevated pTau levels. However, with the integration of cognitive modeling, we were able to predict pTau status with an AUC of 0.71. This suggests that, even though individuals with elevated pTau status did not differ from age-matched controls on traditional metrics, the cognitive mechanisms inferred from cognitive modeling distinguished these individuals. This highlights the nuanced detection capabilities of cognitive modeling, emphasizing its potential in identifying early markers of cognitive decline associated with AD pathology.

### 4.3. Changes in guessing strategies as a function of cognitive impairment, and biomarker status

When applying cognitive modeling to predict AD biomarker and cognitive status, one critical emerging theme was the role of guessing strategies. Guessing strategies, or response biases, change in amnestic patients and individuals with dementia, therefore it is worthwhile to investigate how they change in people at risk for AD. Notably, individuals with increased biomarker levels tended to shift their guessing bias from “similar” to “old” and this shift became more pronounced among those with MCI, suggesting a continuum of change. This shift towards guessing “old” aligns with other work demonstrating that MCI is associated with more liberal response biases on recognition memory tasks (Budson et al., 2000, 2001). One plausible explanation is that individuals build up gist, or low-resolution representations, during a task and, unlike younger adults, cannot rely on item-level detail memory. Thus, without high fidelity memory for details, individuals with MCI may over rely on gist and therefore exhibit a more liberal response bias (Budson et al., 2001; Deason et al., 2012). On the MST, this would cause as a shift towards guessing “old”, which is what we observe. Therefore, this supports that looking at the changes in response biases is important when identifying individuals at risk for AD. Important to note, these changes were more reliably seen when employing cognitive modeling compared to the addition of raw traditional metrics. These outcomes underscore the superiority of cognitive modeling in recognizing individuals at risk of AD, validating its utility as a digital biomarker for early detection of the disease.

### 4.4. Conclusion

Here, we asked if cognitive modeling of the MST could be utilized as a digital biomarker for identifying individuals at risk for AD. We demonstrated that, in addition to predicting memory deficits and MCI, cognitive modeling of the MST could predict both amyloid and pTau status in older adults with AUCs of greater than 0.7 in older adults without signs of cognitive decline.

This suggests that cognitive modeling of the MST holds significant potential as a non-invasive, efficient screening tool within the clinical setting.

## 5. Acknowledgements

We thank Alexandra Trelle, Anthony Wagner, and Elizabeth Mormino for graciously sharing their behavioral and CSF data from the Stanford Aging and Memory Study.

## 8. References

Ally, B. A., Hussey, E. P., Ko, P. C., and Molitor, R. J. (2013). Pattern separation and pattern completion in Alzheimer’s disease: evidence of rapid forgetting in amnestic mild cognitive impairment. Hippocampus 23, 1246–1258. doi: 10.1002/hipo.22162

Bakker, A., Kirwan, C. B., Miller, N. I., and Stark, C. E. L. (2008). Pattern separation in the human hippocampal CA3 and dentate gyrus. Science 319, 1640–1642.

Barthélemy, N. R., Salvadó, G., Schindler, S., He, Y., Janelidze, S., Collij, L. E., et al. (2024). Highly Accurate Blood Test for Alzheimer’s Disease Comparable or Superior to Clinical CSF Tests. Nat. Med. doi: 10.1038/s41591-024-02869-z

Belliart-Guérin, G., and Planche, V. (2023). Mnemonic Discrimination Performance in a Memory Clinic: A Pilot Study. J. Alzheimers Dis. 94, 1527–1534. doi: 10.3233/JAD-230221

Berron, D., Cardenas-Blanco, A., Bittner, D., Metzger, C. D., Spottke, A., Heneka, M. T., et al. (2019). Higher CSF Tau Levels Are Related to Hippocampal Hyperactivity and Object Mnemonic Discrimination in Older Adults. J. Neurosci. 39, 8788–8797. doi: 10.1523/JNEUROSCI.1279-19.2019

Boxer, A. L., and Sperling, R. (2023). Accelerating Alzheimer’s therapeutic development: The past and future of clinical trials. Cell 186, 4757–4772. doi: 10.1016/j.cell.2023.09.023

Budson, A. E., Daffner, K. R., Desikan, R., and Schacter, D. L. (2000). When false recognition is unopposed by true recognition: Gist-based memory distortion in alzheimer’s disease. Neuropsychology 14, 277–287.

Budson, A. E., Desikan, R., Daffner, K. R., and Schacter, D. L. (2001). Perceptual false recognition in alzheimer’s disease. Neuropsychology 15, 230–243.

Chwiesko, C., Janecek, J., Doering, S., Hollearn, M., McMillan, L., Vandekerckhove, J., et al. (2023). Parsing memory and nonmemory contributions to age-related declines in mnemonic discrimination performance: a hierarchical Bayesian diffusion decision modeling approach. Learn. Mem. 30, 296–309. doi: 10.1101/lm.053838.123

Crum, R. M., Anthony, J. C., Bassett, S. S., and Folstein, M. F. (1993). Population-based norms for the Mini-Mental State Examination by age and educational level. JAMA 269, 2386– 91.

Dagum, P. (2018). Digital biomarkers of cognitive function. Npj Digit. Med. 1, 10. doi: 10.1038/s41746-018-0018-4

Deason, R. G., Hussey, E. P., Ally, B. A., and Budson, A. E. (2012). Changes in response bias with different study-test delays: Evidence from young adults, older adults, and patients with Alzheimer’s disease. Neuropsychology 26, 119–126. doi: 10.1037/a0026330

Desikan, R. S., McEvoy, L. K., Thompson, W. K., Holland, D., Brewer, J. B., Aisen, P. S., et al. (2012). Amyloid-β–Associated Clinical Decline Occurs Only in the Presence of Elevated P-tau. Arch. Neurol. 69. doi: 10.1001/archneurol.2011.3354

Ding, Z., Lee, T.-L., and Chan, A. S. (2022). Digital Cognitive Biomarker for Mild Cognitive Impairments and Dementia: A Systematic Review. J. Clin. Med. 11, 4191. doi: 10.3390/jcm11144191

Donohue, M. C., Sperling, R. A., Petersen, R., Sun, C.-K., Weiner, M. W., Aisen, P. S., et al. (2017). Association Between Elevated Brain Amyloid and Subsequent Cognitive Decline Among Cognitively Normal Persons. JAMA 317, 2305. doi: 10.1001/jama.2017.6669

Gefen, T., Peterson, M., Papastefan, S. T., Martersteck, A., Whitney, K., Rademaker, A., et al. (2015). Morphometric and Histologic Substrates of Cingulate Integrity in Elders with Exceptional Memory Capacity. J. Neurosci. 35, 1781–1791. doi: 10.1523/JNEUROSCI.2998-14.2015

Gefen, T., Shaw, E., Whitney, K., Martersteck, A., Stratton, J., Rademaker, A., et al. (2014). Longitudinal Neuropsychological Performance of Cognitive SuperAgers. J. Am. Geriatr. Soc. 62, 1598–1600. doi: 10.1111/jgs.12967

Gellersen, H. M., Trelle, A. N., Henson, R. N., and Simons, J. S. (2021). Executive function and high ambiguity perceptual discrimination contribute to individual differences in mnemonic discrimination in older adults. Cognition 209, 104556. doi: 10.1016/j.cognition.2020.104556

Hansson, O., Blennow, K., Zetterberg, H., and Dage, J. (2023). Blood biomarkers for Alzheimer’s disease in clinical practice and trials. *Nat*. Aging 3, 506–519. doi: 10.1038/s43587-023-00403-3

Harrison, T. M., Weintraub, S., Mesulam, M.-M., and Rogalski, E. (2012). Superior Memory and Higher Cortical Volumes in Unusually Successful Cognitive Aging. J. Int. Neuropsychol. Soc. 18, 1081–1085. doi: 10.1017/S1355617712000847

Hedden, T., Oh, H., Younger, A. P., and Patel, T. A. (2013). Meta-analysis of amyloid-cognition relations in cognitively normal older adults. Neurology 80, 1341–1348. doi: 10.1212/WNL.0b013e31828ab35d

Jack, C. R., Bennett, D. A., Blennow, K., Carrillo, M. C., Dunn, B., Haeberlein, S. B., et al. (2018). NIA-AA Research Framework: Toward a biological definition of Alzheimer’s disease. Alzheimers Dement. J. Alzheimers Assoc. 14, 535–562. doi: 10.1016/j.jalz.2018.02.018

Jia, J., Ning, Y., Chen, M., Wang, S., Yang, H., Li, F., et al. (2024). Biomarker Changes during 20 Years Preceding Alzheimer’s Disease. N. Engl. J. Med. 390, 712–722. doi: 10.1056/NEJMoa2310168

Jutten, R. J., Rentz, D. M., Fu, J. F., Mayblyum, D. V., Amariglio, R. E., Buckley, R. F., et al. (2022). Monthly At-Home Computerized Cognitive Testing to Detect Diminished Practice Effects in Preclinical Alzheimer’s Disease. Front. Aging Neurosci. 13, 800126. doi: 10.3389/fnagi.2021.800126

Kim, S., Adams, J. N., Chappel-Farley, M. G., Keator, D., Janecek, J., Taylor, L., et al. (2023). Examining the diagnostic value of the mnemonic discrimination task for classification of cognitive status and amyloid-beta burden. Neuropsychologia 191, 108727. doi: 10.1016/j.neuropsychologia.2023.108727

Kirwan, C. B., Hartshorn, A., Stark, S. M., Goodrich-Hunsaker, N. J., Hopkins, R. O., and Stark, C. E. (2012). Pattern separation deficits following damage to the hippocampus. Neuropsychologia 50, 2408–14. doi: 10.1016/j.neuropsychologia.2012.06.011

Kirwan, C. B., and Stark, C. E. L. (2007). Overcoming interference: An fMRI investigation of pattern separation in the medial temporal lobe. Learn. Mem. 14, 625–633.

Lacy, J. W., Yassa, M. A., Stark, S. M., Muftuler, L. T., and Stark, C. E. (2011). Distinct pattern separation related transfer functions in human CA3/dentate and CA1 revealed using high-resolution fMRI and variable mnemonic similarity. Learn Mem 18, 15–8. doi: 10.1101/lm.1971111

Lee, M. D., Bock, J. R., Cushman, I., and Shankle, W. R. (2020). An application of multinomial processing tree models and Bayesian methods to understanding memory impairment. J. Math. Psychol. 95, 102328. doi: 10.1016/j.jmp.2020.102328

Lee, M. D., and Stark, C. E. L. (2023). Bayesian modeling of the Mnemonic Similarity Task using multinomial processing trees. Behaviormetrika 50, 517–539. doi: 10.1007/s41237-023-00193-3

Li, Y., Yen, D., Hendrix, R. D., Gordon, B. A., Dlamini, S., Barthélemy, N. R., et al. (2024). Timing of Biomarker Changes in Sporadic Alzheimer’s Disease in Estimated Years from Symptom Onset. Ann. Neurol., ana.26891. doi: 10.1002/ana.26891

Lim, Y. Y., Snyder, P. J., Pietrzak, R. H., Ukiqi, A., Villemagne, V. L., Ames, D., et al. (2016). Sensitivity of composite scores to amyloid burden in preclinical Alzheimer’s disease: Introducing the Z-scores of Attention, Verbal fluency, and Episodic memory for Nondemented older adults composite score. Alzheimers Dement. Diagn. Assess. Dis. Monit. 2, 19–26. doi: 10.1016/j.dadm.2015.11.003

Maass, A., Berron, D., Harrison, T. M., Adams, J. N., La Joie, R., Baker, S., et al. (2019). Alzheimer’s pathology targets distinct memory networks in the ageing brain. Brain 142, 2492–2509. doi: 10.1093/brain/awz154

Macdougall, A., Whitfield, T., Needham, K., Schott, J. M., Frost, C., and Walker, Z. (2024). Predicting progression to Alzheimer’s disease dementia using cognitive measures. Int. J. Geriatr. Psychiatry 39, e6067. doi: 10.1002/gps.6067

McMahon, P. M., Araki, S. S., Sandberg, E. A., Neumann, P. J., and Gazelle, G. S. (2003). Cost-Effectiveness of PET in the Diagnosis of Alzheimer Disease. Radiology 228, 515–522. doi: 10.1148/radiol.2282020915

Mohs, R., Bakker, A., Rosenzweig-Lipson, S., Rosenblum, M., Barton, R. L., Albert, M. S., et al. (2024). The HOPE4MCI study: A randomized double-blind assessment of AGB101 for the treatment of MCI due to AD. Alzheimers Dement. Transl. Res. Clin. Interv. 10, e12446. doi: 10.1002/trc2.12446

Morrison, J. H., and Hof, P. R. (2002). “Chapter 37 Selective vulnerability of corticocortical and hippocampal circuits in aging and Alzheimer’s disease,” in *Progress in Brain Research*, (Elsevier), 467–486. doi: 10.1016/S0079-6123(02)36039-4

Mulhauser, K., Giordani, B., Kavcic, V., May, L. D. N., Bhaumik, A., Shair, S., et al. (2023). Utility of Diffusion Modeling of Cogstate Brief Battery Test Performance in Detecting Mild Cognitive Impairment. Assessment 30, 847–855. doi: 10.1177/10731911211069089

Norman, K. A., Detre, G., and Polyn, S. M. (2001). “Computational Models of Episodic Memory,” in The Cambridge Handbook of Computational Psychology, ed. R. Sun (Cambridge University Press), 189–225. doi: 10.1017/CBO9780511816772.011

Ossenkoppele, R., Pichet Binette, A., Groot, C., Smith, R., Strandberg, O., Palmqvist, S., et al. (2022). Amyloid and tau PET-positive cognitively unimpaired individuals are at high risk for future cognitive decline. Nat. Med. 28, 2381–2387. doi: 10.1038/s41591-022-02049-x

Papp, K. V., Jutten, R. J., Soberanes, D., Weizenbaum, E., Hsieh, S., Molinare, C., et al. (2023). Early Detection of Amyloid-Related Changes in Memory among Cognitively Unimpaired Older Adults with Daily Digital Testing. Ann. Neurol., ana.26833. doi: 10.1002/ana.26833

Papp, K. V., Rentz, D. M., Maruff, P., Sun, C.-K., Raman, R., Donohue, M. C., et al. (2020). The Computerized Cognitive Composite (C3) in A4, an Alzheimer’s Disease Secondary Prevention Trial. J. Prev. Alzheimers Dis., 1–9. doi: 10.14283/jpad.2020.38

Radhakrishnan, H., Bennett, I. J., and Stark, C. E. (2022). Higher-order multi-shell diffusion measures complement tensor metrics and volume in gray matter when predicting age and cognition. NeuroImage 253, 119063. doi: 10.1016/j.neuroimage.2022.119063

Raftery, A. E. (1995). Bayesian Model Selection in Social Research. Sociol. Methodol. 25, 111. doi: 10.2307/271063

Rey, A. (1941). L’examen psychologique dans les cas d’encephalopathie traumatique. Arch Psychol 28, 286–340.

Sabuncu, M. R. (2011). The Dynamics of Cortical and Hippocampal Atrophy in Alzheimer Disease. Arch. Neurol. 68, 1040. doi: 10.1001/archneurol.2011.167

Seabold, S., and Perktold, J. (2010). Statsmodels: Econometric and Statistical Modeling with Python. 5.

Small, S. A., Perera, G. M., DeLaPaz, R., Mayeux, R., and Stern, Y. (1999). Differential regional dysfunction of the hippocampal formation among elderly with memory decline and Alzheimer’s disease. Ann. Neurol. 45, 466–472.

Small, S. A., Schobel, S. A., Buxton, R. B., Witter, M. P., and Barnes, C. A. (2011). A pathophysiological framework of hippocampal dysfunction in ageing and disease. Nat. Rev. Neurosci. 12, 585–601. doi: 10.1038/nrn3085

Sperling, R. A., Aisen, P. S., Beckett, L. A., Bennett, D. A., Craft, S., Fagan, A. M., et al. (2011). Toward defining the preclinical stages of Alzheimer’s disease: Recommendations from the National Institute on Aging-Alzheimer’s Association workgroups on diagnostic guidelines for Alzheimer’s disease. Alzheimers Dement. 7, 280–292. doi: 10.1016/j.jalz.2011.03.003

Sperling, R. A., Karlawish, J., and Johnson, K. A. (2013). Preclinical Alzheimer disease—the challenges ahead. Nat. Rev. Neurol. 9, 54–58. doi: 10.1038/nrneurol.2012.241

Stark, S. M., Kirwan, C. B., and Stark, C. E. L. (2019). Mnemonic Similarity Task: A Tool for Assessing Hippocampal Integrity. Trends Cogn. Sci. 23, 938–951. doi: 10.1016/j.tics.2019.08.003

Stark, S. M., Yassa, M. A., Lacy, J. W., and Stark, C. E. (2013). A task to assess behavioral pattern separation (BPS) in humans: Data from healthy aging and mild cognitive impairment. Neuropsychologia 51, 2442–9. doi: 10.1016/j.neuropsychologia.2012.12.014

Student (1908). The Probable Error of a Mean. Biometrika 6, 1–25. doi: 10.2307/2331554

Tombaugh, T. N. (2004). Trail Making Test A and B: normative data stratified by age and education. Arch. Clin. Neuropsychol. Off. J. Natl. Acad. Neuropsychol. 19, 203–214. doi: 10.1016/S0887-6177(03)00039-8

Trelle, A. N., Carr, V. A., Wilson, E. N., Swarovski, M. S., Hunt, M. P., Toueg, T. N., et al. (2021). Association of CSF Biomarkers With Hippocampal-Dependent Memory in Preclinical Alzheimer Disease. Neurology 96. doi: 10.1212/WNL.0000000000011477

Wagenmakers, E.-J., and Farrell, S. (2004). AIC model selection using Akaike weights. Psychon Bull. Rev. 11, 192–196. doi: 10.3758/BF03206482

Wang, W., Peng, J., Hou, J., Yuan, Z., Xie, W., Mao, G., et al. (2023). Predicting mild cognitive impairment progression to Alzheimer’s disease based on machine learning analysis of cortical morphological features. Aging Clin. Exp. Res. 35, 1721–1730. doi: 10.1007/s40520-023-02456-1

Wechsler, D. (1997a). Wechsler Adult Instelligence Scale (WAIS-III): Administration and scoring manual. San Antonio, TX: The Psychological Corporation.

Wechsler, D. (1997b). Wechsler Adult Intelligence Scale (WAIS-III): Administration and scoring manual. San Antonio, TX: The Psychological Corporation.

Wechsler, D. (1997c). Wechsler Memory Scale - Third Edition (WMS-III). San Antonio, TX: The Psychological Corporation.

Wittenberg, R., Knapp, M., Hu, B., Comas-Herrera, A., King, D., Rehill, A., et al. (2019). The costs of dementia in England. Int. J. Geriatr. Psychiatry 34, 1095–1103. doi: 10.1002/gps.5113

Yassa, M. A., Stark, S. M., Bakker, A., Albert, M. S., Gallagher, M., and Stark, C. E. L. (2010). High-resoltuion structural and functional MRI of hippocampal CA3 and dentate gyrus in patients with amnestic mild cognitive impairment. NeuroImage 51, 1242–1252.

